# Guess Who? Identifying individuals from their brain natural frequency fingerprints

**DOI:** 10.1101/2025.01.21.634078

**Authors:** Lydia Arana, Juan José Herrera-Morueco, Javier Santonja, Almudena Capilla

## Abstract

Neural oscillations are critical for brain function and cognition. Thus, identifying the typical or natural oscillatory frequencies of the brain is an important first step for understanding its functional architecture. Recently, a data-driven algorithm has been developed for mapping the brain’s natural frequencies throughout the whole cortex, free of anatomical and frequency-band constraints. However, an important limitation of this methodology is that it yields robust results only at the group level. Here, we aimed to adapt this algorithm to improve the quality of the single-subject maps of natural frequencies obtained from magnetoencephalography (MEG) recordings. To achieve this goal, we incorporated two modifications to the original method: (1) increasing the number of individual power spectra to be assigned to each k-means cluster, and (2) smoothing across neighboring voxels. To assess the quality of the single-subject maps, we relied on the fingerprinting technique. Our results show a high degree of accuracy in individual identification, both within a single recording session and across separate sessions. Furthermore, we were able to identify individuals by their natural frequency fingerprints, even with a gap of over four years between sessions. This demonstrates the robustness of the single-subject mapping of natural frequencies and opens new opportunities for identification of pathological variations in intrinsic oscillatory activity in individual subjects.

## 1. Introduction

Electromagnetic oscillations are spontaneously generated by the brain and are postulated to play a key role in neural communication, ultimately supporting the mechanisms of cognition (Buzsáki & Watson, 2012; Fries, 2015). Gaining a deeper understanding of neural oscillations is therefore essential for deciphering the brain’s communication code and for unraveling how cognitive processes emerge from the coordinated activity of neural networks. For this purpose, an important initial step is to identify which are the natural or typical oscillatory frequencies in different parts of the brain. This has been explored through two main strategies: directly perturbing intrinsic oscillations by a single pulse of transcranial magnetic stimulation (TMS) (Amengual et al., 2019; Ferrarelli et al., 2012; Rosanova et al., 2009) or characterizing region-specific spectral features from electro-/magneto-encephalography (EEG/MEG) and intracranial EEG (iEEG) recordings (Capilla et al., 2022; Frauscher et al., 2018; Kalamangalam et al., 2020; Keitel & Gross, 2016; Mahjoory et al., 2020; Mellem et al., 2017). Interestingly, spectral profiles are so consistent that they can be used to differentiate and classify brain regions based on their intrinsic oscillatory activity (Keitel & Gross, 2016; Komorowski et al., 2023).

With the aim of providing a detailed atlas of the natural frequencies of the healthy human brain at rest, Capilla and colleagues (2022) developed a novel data-driven algorithm for mapping natural frequencies on a voxel-by-voxel basis, free of anatomical and frequency-band constraints. The shift in focus from the conventional frequency bands to the study of individual frequencies allowed them to identify, for example, different generators of oscillatory activity in sensory regions (visual, auditory and tactile), all of them characterized by oscillations within the alpha band, but each with a distinctive frequency (e.g., ∼12 Hz in visual areas, ∼8 Hz in auditory areas). Overall, ongoing oscillations showed a region-specific organization, which was structured along two gradients of increasing frequency, from medial to lateral and from posterior to anterior brain areas. In particular, medial frontal and temporal regions were characterized by slow oscillations (delta and theta, 0.5-4 Hz and 4-8 Hz, respectively), posterior occipito-temporal cortices were characterized by alpha-band oscillations (8-13 Hz), while lateral frontal and parietal cortices exhibited oscillations in the beta range (13-30 Hz), in line with previous reports (Frauscher et al., 2018; Keitel & Gross, 2016; Mellem et al., 2017; Niso et al., 2019).

However, an important limitation of this methodology is that it only provides robust results at the group level. Although individual brain patterns of natural frequencies can, in principle, be computed, they are of modest quality. Still, obtaining single-subject maps is essential for statistical purposes as well as to identify individual normal/pathological variations. Accordingly, in this study we aimed to adapt Capilla et al.’s approach to improve the quality of the single-subject mapping of the brain’s natural frequencies.

To determine whether brain maps of natural frequencies are robust at the single-subject level, we relied on the fingerprinting technique. The concept of brain fingerprinting has gained traction in recent years within the neuroscientific literature, inspired by traditional forensic and biometric practices, where fingerprints are used for individual identification. Several neuroimaging-based approaches have been proposed to extract unique features of brain activity. The most established procedure makes use of functional magnetic resonance imaging (fMRI) to obtain functional connectivity profiles, which are then employed as fingerprints to identify a target subject among others (Amico & Goñi, 2018; Finn et al., 2015; Mantwill et al., 2022; Miranda-Dominguez et al., 2014; Van De Ville et al., 2021). Only recently, more complex spectral and electrophysiological connectivity profiles have also been extracted from MEG recordings to enable individual differentiation (Colenbier et al., 2023; Da Silva Castanheira et al., 2021, 2024; Sareen et al., 2021). In both cases, the number of features employed is considerably large. Here, we propose using a relatively simple index of brain function, the brain pattern of natural frequencies at rest (a vector with 1925 elements, i.e., each voxels’ natural frequency) as a brain fingerprint to identify individuals.

## 2. Material and methods

### 2.1 Participants and data acquisition

All procedures complied with the Declaration of Helsinki and were approved by the institutional ethics committees. Data were obtained from The Open MEG Archive (OMEGA; Niso et al., 2016), an open-access database which provides anonymized resting-state MEG recordings and T1-weighted Magnetic Resonance Images (MRIs). Brain activity was recorded in a magnetically shielded room at the Montreal Neurological Institute (MNI, McGill University) using a whole-head CTF MEG system with 275 axial gradiometers and 26 reference sensors at a 2400 Hz sampling rate. An anti-aliasing low-pass filter at 600 Hz was applied online, as well as CTF third-order gradient compensation. Participants remained awake with eyes open on a fixation cross for 5 minutes.

#### Within-session group

The within-session group included 128 healthy volunteers (68 males, 118 right-handed, 30.5 ± 12.4 [M ± SD] years, ranging from 19 to 73 years) with one session per participant. This sample was used to conduct fingerprinting using data from both halves of a single session for each participant (Fig. 1a).

**Figure 1.**
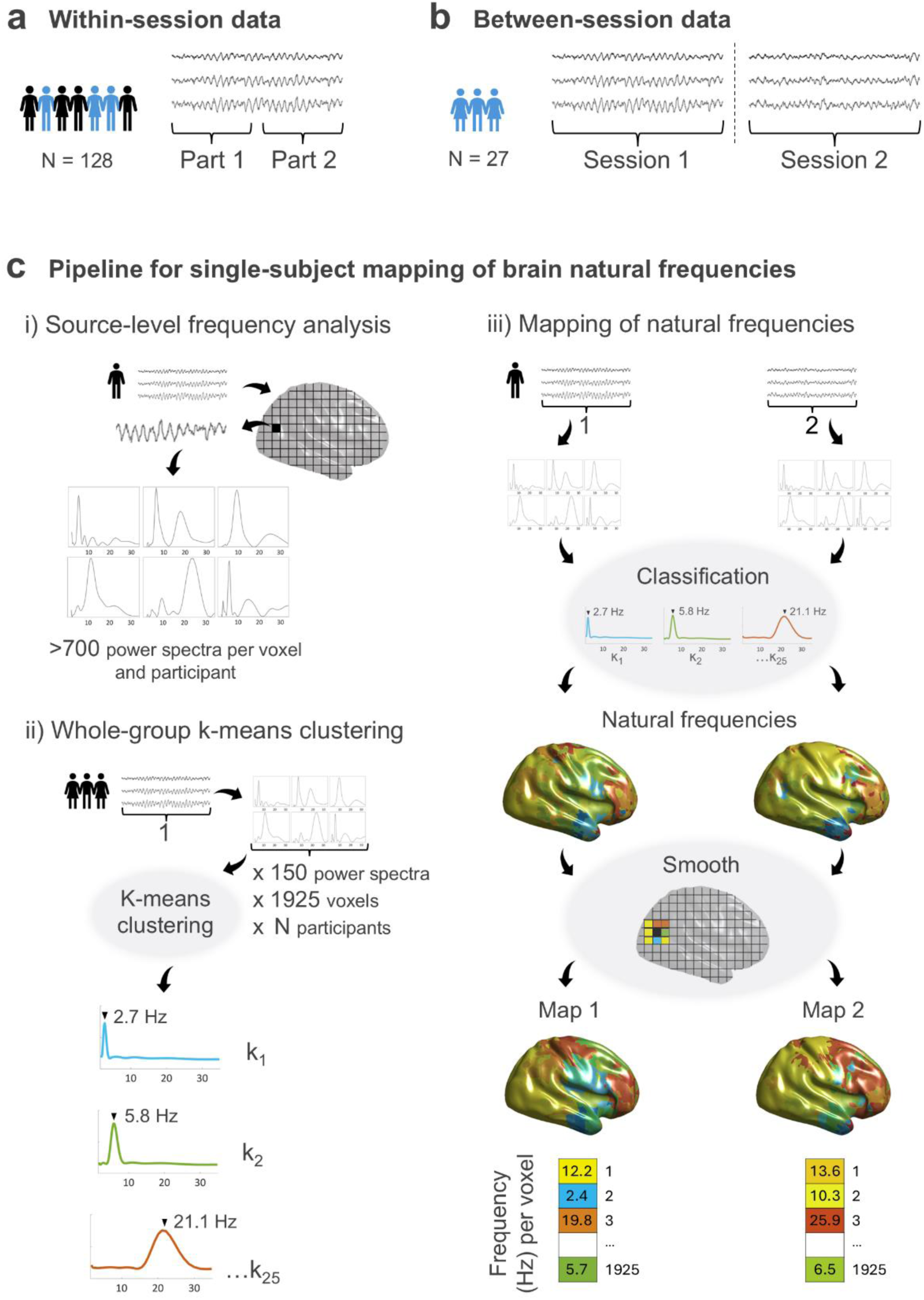
MEG data and pipeline for obtaining individual brain maps of natural frequencies. **(a)** Within-session data. A single session from each participant was divided into two parts and pre-processed separately. **(b)** Between-session data. The two MEG sessions recorded at different times for each participant were pre-processed separately. **(c)** Pipeline for mapping the brain’s natural frequencies. i) Source reconstruction and frequency analysis of each part/session. ii) Whole-group k-means clustering was conducted on 150 spectra per voxel and participant from Part/Session 1. iii) Brain mapping of single-subject natural frequencies was derived separately for Part/Session 1 and Part/Session 2. First, single-subject power spectra obtained from one instance of the data were classified into the whole-group clusters. Next, the peak of the most typical power spectrum defined the natural frequency of each voxel, which were smoothed across neighboring voxels. Final brain maps for Part/Session 1 and Part/Session 2 were composed of single vectors of 1925 natural frequencies, i.e. one value per voxel.

#### Between-session group

The between-session group comprised a subset of 27 individuals from the whole sample with more than one MEG session on the same or on different days (18 males, 26 right-handed, 27.5 ± 6.0 years, age range 21-46 years in the first session; time between sessions: 299 ± 411 days, ranging from 0 to 1696 days). This subsample was intended to assess the performance of the fingerprinting approach on a subsequent recording (Fig. 1b). When more than one additional session was available, we discarded those containing more artifacts.

### 2.2 Pre-processing and reconstruction of source-level activity

Every recording was initially trimmed to 5 minutes to ensure equal duration for all participants. For the within-session group, the recording was split into two halves of 2.5 minutes each; from here on, they will be referred to as Part 1 and Part 2. For the between-session group, both sessions remained as 5-minute recordings; we will refer to them as Session 1 and Session 2. Each part/session was analyzed separately, as described in the following paragraphs.

The analysis pipeline for data pre-processing and source reconstruction was identical to that described in Capilla et al. (2022). Analyses were carried out using FieldTrip (version 20230118; Oostenveld et al., 2011) and in-house Matlab code. The scripts necessary to reproduce all the analysis and figures are available at https://github.com/necog-UAM.

The MEG signal was denoised using Principal Component Analysis (PCA). Then, data were high-pass filtered at 0.05 Hz with a third-order Butterworth filter. The power line artifact at 60 Hz (and harmonics at 120 and 180 Hz) was reduced by means of spectrum interpolation (Leske & Dalal, 2019). The MEG signal was then demeaned, detrended, and resampled at 512 Hz. Artifacts were corrected with Independent Component Analysis (ICA), after a PCA dimensionality reduction to 40 components. Any data segments showing remaining artifacts upon visual inspection were excluded from further analysis.

Individual T1-weighted MRIs were co-registered to the MEG coordinate system using a semi-automatic procedure based on a modified version of the Iterative Closest Point algorithm (Besl & McKay, 1992). The forward model was computed using a realistic single shell volume conductor model (Nolte, 2003). A standard MNI grid with 1-cm resolution was adapted to each individual’s brain volume and lead fields were computed for each grid point. Voxels in regions outside the cerebral cortex or the hippocampus (i.e., cerebellum and subcortical structures) were excluded, leaving 1925 voxels for further analysis.

To reconstruct source-level time series, we employed linearly constrained minimum variance (LCMV) beamforming (Van Veen et al., 1997). The spatial filter weights were derived from the covariance of the artifact-free data, with the regularization parameter lambda set to 10%. These beamforming weights were then used to estimate source-space time series from the sensor-level data.

### 2.3 Frequency analysis on source-level data

We applied a Hanning-tapered sliding window Fourier transform in steps of 200 ms on the source-reconstructed MEG signal. Spectral analysis was performed at a higher temporal resolution compared to Capilla et al. (2022) (i.e., every 200 ms instead of 500 ms) to increase the number of power spectra and thereby enhance the quality of the single-subject maps of natural frequencies (see Fig. 1c, i). Power was computed for 61 frequency bins logarithmically spaced from 1.7 to 34.5 Hz, as we previously observed that higher frequencies reflected artifactual activity (Capilla et al., 2022). The width of the sliding window was frequency-dependent and adapted to a length of 5 cycles per frequency bin to attenuate the 1/f aperiodic component. We discarded 5.8-s time intervals (i.e., 10 cycles at the lowest frequency of 1.7 Hz) at the beginning and the end of every artifact-free data segment to avoid edge effects. Thus, for the within-session group, we obtained a set of 778 ± 29 power spectra per voxel and participant from Part 1 and 783 ± 28 spectra from Part 2. For the between-session group, we obtained 1286 ± 20 spectra from Session 1 and 1259 ± 19 spectra from Session 2. Finally, to account for the center of the head bias, each spectrum was expressed as relative power by dividing the power at each frequency by the absolute power summed over the whole spectrum.

### 2.4 Whole-group cluster analysis of power spectra

To identify different patterns of source-reconstructed oscillatory activity, we performed a whole-group k-means clustering with the cosine as the distance metric to maximize differences between clusters based on the shape of the spectra (Keitel & Gross, 2016). Clustering was performed separately for the within-session and the between-session groups, randomly selecting 150 power spectra per voxel and participant from Part 1 and Session 1, respectively. We trained the k-means clustering model only with data from Part/Session 1 to avoid biasing the identification of data from Part/Session 2. Thus, we introduced a total of 150 x 1925 x 128 power spectra for the within-session group, and 150 x 1925 x 27 power spectra for the between-session group. The number of clusters was set to 25. To achieve optimal results, we run 5 clustering replicates with a maximum of 200 iterations and selected the solution with the lowest sum of distances (see Capilla et al., 2022) (Fig. 1c, ii).

### 2.5 Single-subject brain maps of natural frequencies

Once the k-means clustering model was trained using data from Part/Session 1 of the entire group, we computed the single-subject maps of natural frequencies. For each individual, we obtained two maps (Part 1 and Part 2 maps for the within-session group participants; Session 1 and Session 2 maps for the between-session group participants), following the procedure below (see Fig. 1c, iii).

For each participant and voxel, we assigned the previously computed power spectra (see section 2.3) to each cluster, by computing the distance between the power spectra and the cluster centroids. Each power spectrum was assigned to the cluster with the smallest distance to its centroid. In addition, to identify the oscillatory frequency characterizing each cluster, we detected the frequency peak of each centroid. Cluster centroids with no peaks were disregarded. In case two peaks were detected (e.g., mu-rhythm), each frequency peak (e.g., 10 and 20 Hz) was weighted at 50%. If more than two peaks were detected, only the two largest were considered.

The first step towards improving the quality of the single-subject maps was to classify a larger number of power spectra per participant (ranging from 428 to 1414 for participants of the within-session group and 1042 to 1364 for participants of the between-session group Vs. 150 spectra used in the original formulation of the method). The proportion of power spectra classified within each cluster was then calculated and normalized across voxels (z-score). We interpolated centroid peak frequencies and their corresponding z-values 10 times to obtain a wider range of frequency values. For each voxel, the oscillatory frequency with the highest z-value was defined as its natural frequency.

Finally, the second measure to improve individual maps was to smooth across neighboring voxels. Since there might be abrupt transitions between the natural frequencies of surrounding voxels (e.g., delta and high-beta frequencies in frontal regions), standard smoothing is not appropriate for these data, and we opted for an alternative approach. For each voxel, we identified its neighbors in a sphere of < 1.5 cm radius. We then computed a t-test against zero across surrounding voxels for each frequency bin and selected the frequency with the highest t-value as the voxel’s natural frequency (i.e., the most representative frequency value within the local vicinity). If the t-value was not statistically significant (p > 0.05) (i.e., the natural frequency was not stable in its neighboring set of voxels), we assigned a missing value to the voxel.

### 2.6 Identification of participants

After obtaining the natural frequency brain maps for every participant in each part/session, we carried out the identification of individual brains, separately for the within-session and the between-session groups. The identification procedure followed the common fingerprinting approach, which is based on the correlation between the first and the second instances of all participants’ data (Amico & Goñi, 2018; Da Silva Castanheira et al., 2021; Finn et al., 2015).

We calculated the Kendall’s tau (τ) correlation coefficients between the Part/Session 1 natural frequency maps and Part/Session 2 maps for all individuals, thus obtaining a square correlation matrix of size participants x participants (128 x 128 in the within-session group, and 27 x 27 in the between-session group). We chose Kendall’s correlation coefficient because our data did not conform to a normal distribution, as assumed by Pearson’s correlation (Croux & Dehon, 2010). We performed a Kolmogorov–Smirnov test to check whether the natural frequencies of each map were normally distributed and found that none of the maps were Gaussian (p < .001).

The diagonal of the correlation matrix represents the autocorrelation of each individual with themselves. The fingerprinting process consists of a search through the rows/columns of the correlation matrix, where the highest correlation coefficient indicates the matching participant and receives a score of 1. The rest of participants in the row/column are thus non-matches and are assigned a score of 0. This process resulted in a final matrix of matches, derived from the two instances of natural frequency maps.

Correct matches or “target matches” occur when participants correctly match with themselves, while “non-target matches” refer to cases where the first map of one participant matches the second map of a different participant. The overall accuracy of the brain fingerprinting was determined by calculating the percentage of correctly identified individuals or target matches.

For descriptive purposes, we calculated the means and standard deviations of the correlation coefficients (τ). Specifically, we computed the descriptive statistics for the self-correlations (the diagonal of the correlation matrix) and for the correlations with others (values outside the diagonal), in absolute values, separately for the within-session and the between-session groups. To test whether self-correlations were statistically higher than correlations with others, we performed paired sample t-tests.

To compute the confidence interval (CI) of the identification accuracy for each group, we employed a bootstrapping procedure, repeated 1000 times. In each iteration, we randomly resampled participants with replacement to create a new dataset of the same size as the original. For each resampled dataset, we repeated the fingerprinting process to obtain an identification accuracy score. This resulted in a distribution of accuracy scores, from which we derived the 95% CI as the interval between the 2.5^th^ and 97.5^th^ percentiles of the distribution.

We also performed a non-parametric permutation test to assess the statistical significance of the fingerprinting accuracy. This consisted of randomly changing (1000 times) the labels of the participants in the second map, for each group separately, and repeating the identification process. We then computed the highest success rate among all iterations.

### 2.7 Identifiability and differentiability of participants

For each group and participant, we calculated both identifiability and differentiability metrics. Identifiability (I_self_; Amico and Goñi, 2018) is used to quantify the reliability of an individual’s identification among the rest of the cohort. Thus, we computed the correlation of each participant with himself/herself (i.e, self-correlation; τ_self_) minus the average of the correlation with the rest of participants (i.e., others-correlation; μ_others_) (I_self_ = τ_self_ − μ_others_). Thus, a high identifiability value would indicate that a participant is easily identifiable among the others (i.e., individuals with a high self-correlation and/or a low correlation with others).

Da Silva Castanheira et al. (2021) extended this notion with the introduction of a normalized measure, the differentiability. Differentiability (D_self_) is defined as the correlation of one participant with himself/herself (i.e., self-correlation; τ_self_) minus the mean of the correlations with others (i.e., others-correlation; μ_others_), divided by the standard deviation of others-correlations (σ_others_) (D_self_ = [τ_self_ − μ_others_] / σ_others_). As in the case of identifiability, high differentiability values would be indicative of individuals with unique or salient brain natural frequency fingerprints, with the advantage of being a normalized value (z-score) which allows for more direct comparisons.

For descriptive purposes, we also calculated the means and standard deviations of identifiability and differentiability for both groups and the means and standard deviations of target matches and non-target matches. Additionally, we conducted two-sample t-tests to assess the hypothesis that identifiability and differentiability were higher for target than for non-target matches.

### 2.8 Brain fingerprinting across time

Finally, we set out to investigate whether the accuracy of the brain natural frequency fingerprinting decreases as the time between the recording sessions increases. We thus focused on the between-session group, as it included two MEG sessions acquired at different points in time, with an interval from 0 to 1696 days (4.6 years).

For this purpose, we conducted two types of analysis. First, we performed a Pearson’s linear correlation between the number of days between sessions and the value of Kendall’s τ for self-correlations, to determine whether self-correlation decreases with increasing time. Then, we statistically tested whether elapsed time was different for target and non-target matches. Since sample size was rather unbalanced for correct compared to incorrect matches (a higher number of correct matches), we applied a Welch’s t-test (Delacre et al., 2017; Derrick & White, 2016).

## 3. Results

### 3.1 Individual brains can be identified from their natural frequency fingerprints within the same session

In this study we aimed to test whether it is possible to obtain robust brain maps of natural frequencies at the single-subject level. Thus, we first assessed if we could identify individual brains from natural frequency maps obtained within the same session in a sample of 128 participants (within-session group).

The Kendall’s τ correlation matrix of the within-session group is depicted in Figure 2a and led to the identification of participants in the subsequent step. As shown in Figure 2b, we correctly identified 119 out of 128 participants or, in other words, the correlation with oneself was higher than with any other participant in 119/128 cases. Hence, the accuracy of the natural frequency fingerprinting within sessions was 92.97% (95% CI [89.45, 98.44]). To determine whether the observed identifications could have been driven by random similarities in correlations across participants, we performed a non-parametric permutation test in which we shuffled the identity of the participants. Across 1000 iterations, the highest success rate achieved was 1/128 (0.78%). Therefore, the probability of obtaining at least 119/128 correct identifications by chance was below .001.

**Figure 2.**
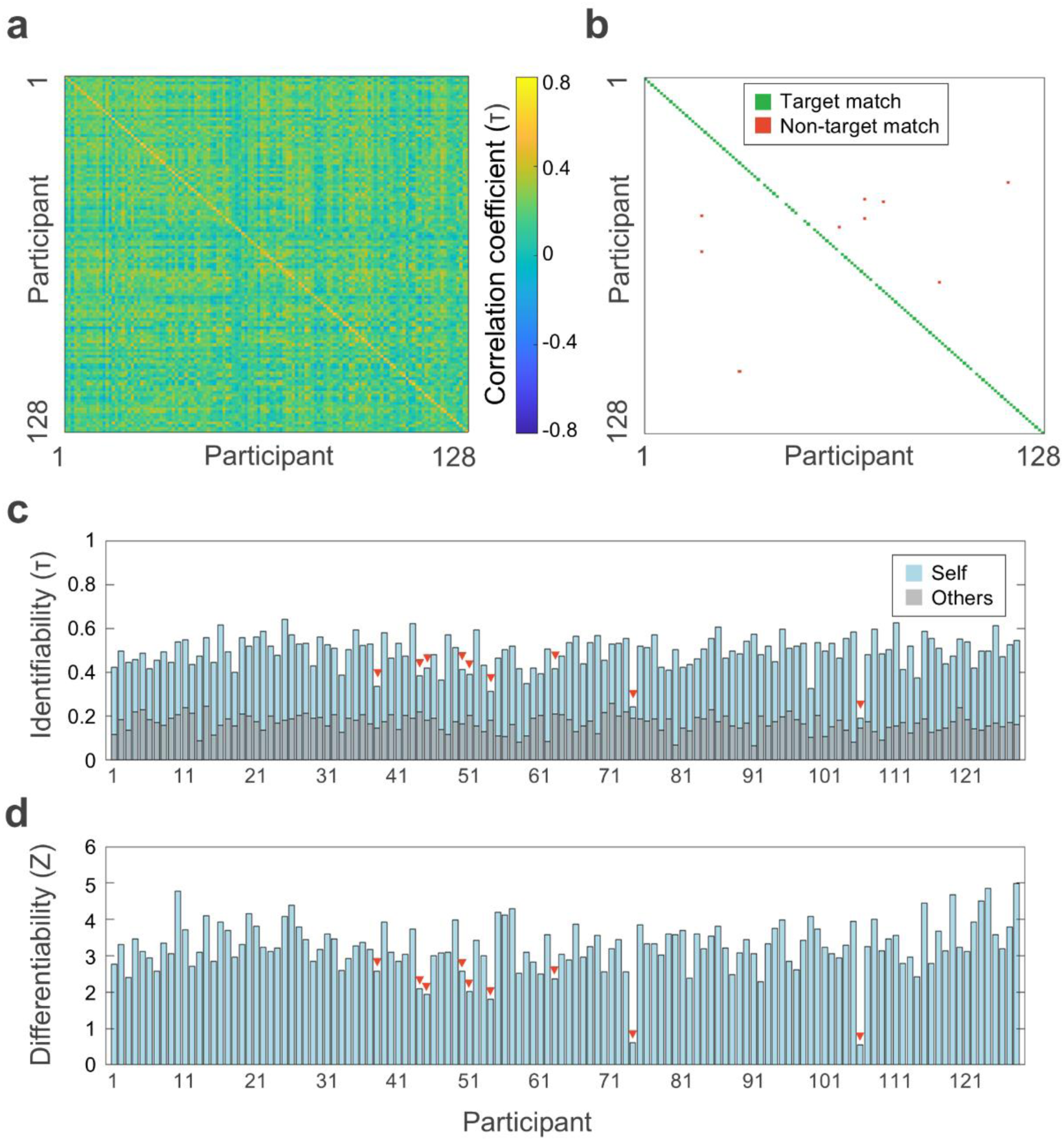
Within-session fingerprinting of natural frequencies. **(a)** Correlation matrix between the natural frequency maps obtained from the two parts of the MEG recording for each participant of the within-session group (N = 128). **(b)** Matrix of correct/incorrect matches. Target matches are represented in green; non-target matches, in red. The graph shows that 119/128 individuals were correctly identified. **(c)** Identifiability of participants (Kendall’s τ). Self-correlation of each participant is depicted in light blue; others-correlation (i.e., mean correlation of each participant with all other participants) is displayed in gray. Red triangles point to the participants that did not match themselves. **(d)** Differentiability provides a normative metric (z-score) of the extent to which the participant’s natural frequency map is distinct from other participants’ maps.

In addition, we found that the Kendall’s τ correlation was higher for self-correlations (0.49 ± 0.08) compared to correlations with others (0.18 ± 0.03) (t_(127)_ = 46.72, p < .001). It is important to note that the interpretation of Kendall’s τ correlation coefficients might differ from the more common Pearson’s r. Although both coefficients range from −1.0 to +1.0, the absolute value of Kendall’s τ is typically about 1.5 times smaller than Pearson’s r (El-Hashash & Shiekh, 2022). Thus, the self-correlation values obtained here would indicate moderate to high correlations.

Figure 2c shows the within-session identifiability of each participant. The overall identifiability of the within-session group was 0.32 ± 0.08, although it was significantly higher for target matches (0.34 ± 0.07) compared to non-target matches (0.16 ± 0.07) (t_(126)_ = 7.26, p < .001).

The z-values for differentiability followed a similar pattern (Fig. 2d). Although the average differentiability of the within-session group was 3.26 ± 0.69, z-values were significantly higher for target matches (3.37 ± 0.56) than for non-target matches (1.84 ± 0.76) (t_(126)_ = 7.73, p < .001).

### 3.2 Individual brains can be identified from their natural frequency fingerprints obtained in separate sessions

Having proved that natural frequency maps can be used to identify individuals with data from the same session, we aimed to test whether we could also identify individuals with data from separate sessions. Our sample consisted of 27 participants who underwent two MEG sessions with an interval of 0 to 1696 days (between-session group).

Figure 3a shows the Kendall’s τ correlation matrix of the between-session group. In this case, 21 out of 27 participants were correctly identified (Fig. 3b) and, therefore, the accuracy of the natural frequency fingerprinting between sessions was 77.78% (95% CI [62.96, 92.59]). The non-parametric permutation test revealed that, if the identity of the participants was randomly shuffled, the highest match rate would be 2/27 (7.41%). Consequently, the accuracy obtained in the between-session group was also above chance level (p < .001).

**Figure 3.**
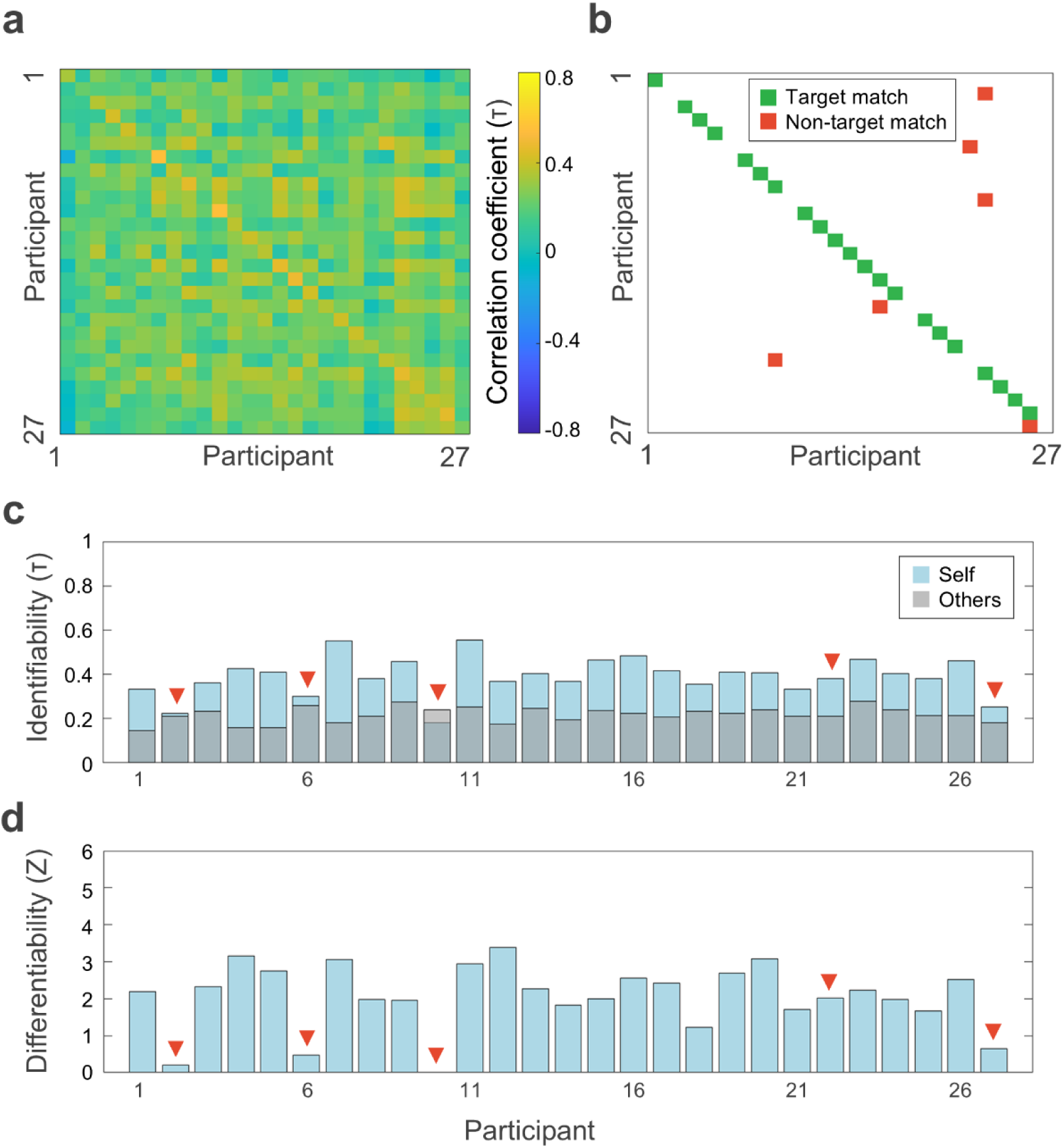
Between-session fingerprinting of natural frequencies. **(a)** Correlation matrix between the natural frequency maps obtained from two separate MEG sessions for each participant of the between-session group (N = 27). **(b)** Matrix of correct/incorrect matches. Target matches are represented in green; non-target matches, in red. The graph shows that 21/27 individuals were correctly identified. **(c)** Identifiability of participants (Kendall’s τ). Self-correlation of each participant is depicted in light blue; others-correlation (i.e., mean correlation of each participant with all other participants) is displayed in gray. Red triangles point to the participants that did not match themselves. **(d)** Differentiability provides a normative metric (z-score) of the extent to which the participant’s natural frequency map is distinct from other participants’ maps.

Similar to the within-session results, we also found that the Kendall’s τ correlation was higher for self-correlations (0.39 ± 0.09) compared to correlations with others (0.22 ± 0.03) (t_(26)_ = 10.19, p < .001) when MEG data comes from two different sessions.

The average between-session identifiability was 0.17 ± 0.09 (Fig. 3c). As for the within-session group results, we also found that identifiability was higher for target matches (0.21 ± 0.06) compared to non-target matches (0.06 ± 0.08) (t_(25)_ = 4.93, p < .001).

Similarly, overall differentiability for the between-session group was 2.03 ± 0.94 (Fig. 3d). Differentiability was also higher for target matches 2.41 ± 0.51 than for non-target matches (0.67 ± 0.87) (t_(25)_ = 6.30, p < .001).

### 3.3 Natural frequency fingerprinting is robust against time

Data from the between-session group allowed us to explore whether participants’ identification decay over time. First, we conducted a Pearson’s linear correlation between elapsed time and self-correlations. This correlation did not result statistically significant (r_(25)_ = - .010, p = .96), indicating that identifiability does not decrease as time between sessions increases (Fig. 4a).

**Figure 4.**
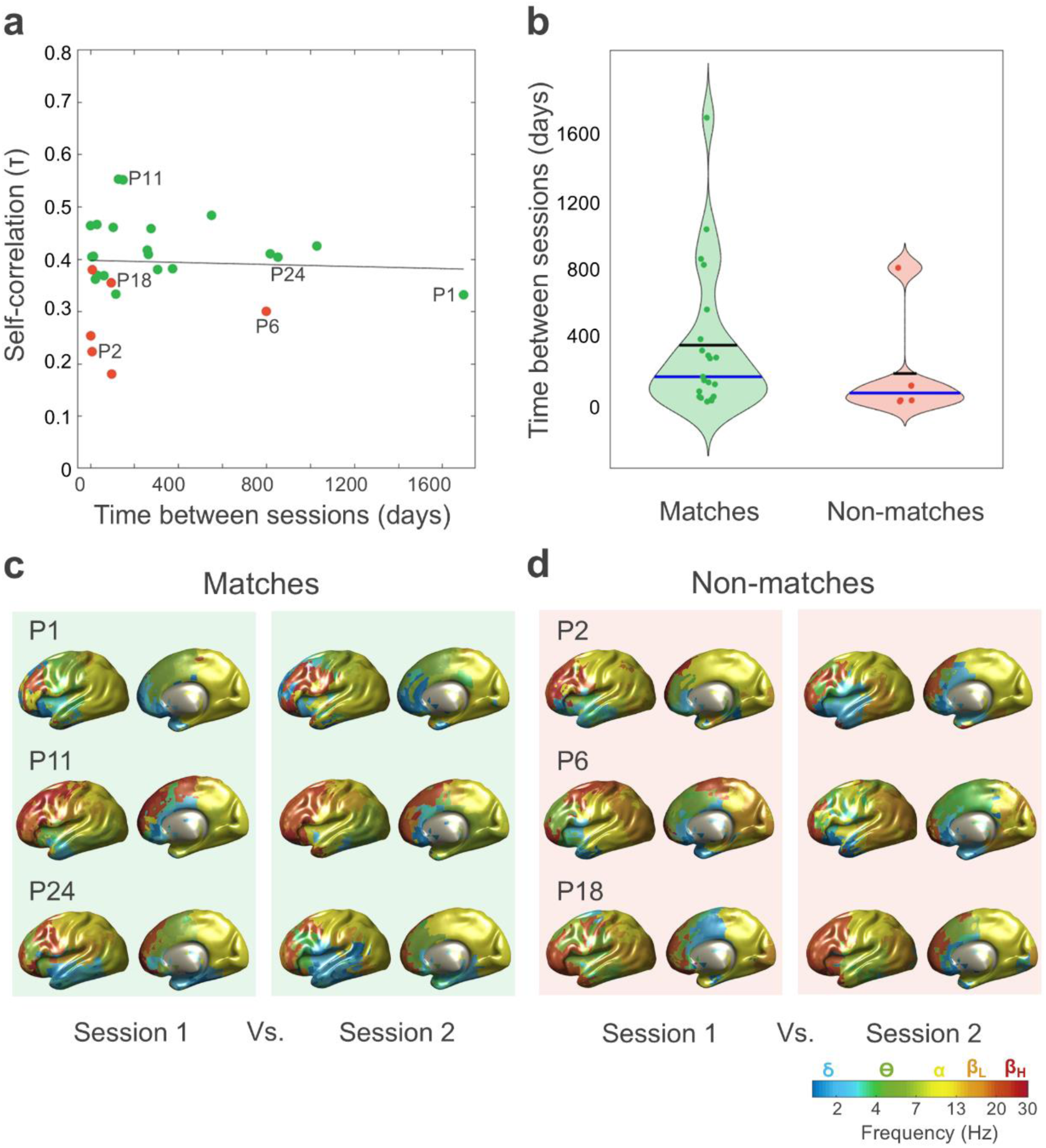
Fingerprinting of natural frequencies is not affected by the interval between sessions. **(a)** Absence of a significant correlation between self-correlation Kendall’s τ coefficients and days elapsed between sessions. Green indicates participants whose first session matched their second, while red indicates those whose sessions did not match. The trend line is shown in black. The letter P (participant) and a number identifies specific individuals, which are further described in the next panels. **(b)** Absence of significant differences in the time elapsed between sessions for correct matches (in green) Vs. incorrect matches (in red). Means are represented by black horizontal lines; blue lines represent the medians. **(c)** Examples of correct matches. Natural frequency brain maps for the two sessions of participants P1, P11, and P24 (>4 years, >4 months, >2 years elapsed between sessions, respectively). **(d)** Examples of incorrect matches. Brain maps of participants P2, P6, and P18 (7 days, >2 years, >3 months elapsed between sessions, respectively). For simplicity, only the lateral view of the left hemisphere and the medial view of the right hemisphere are shown.

We then tested whether elapsed time was different for target and non-target matches. We found that the number of days between sessions did not significantly differ between target matches (337 ± 434 days) and non-target matches (167 ± 312 days) (t_(11.17)_ = 1.07, p = .31, 95% CI [-180, 518]) (Fig. 4b). Taken together, these results suggest that the time elapsed between recording sessions does not influence identification accuracy and, therefore, natural frequency fingerprinting is robust against time.

Figure 4c shows some representative brain maps of natural frequencies obtained from the two separate sessions of correctly identified participants. Note the similarity between both maps even in cases where MEG recordings are several years apart (e.g., participant 1 or 24). Finally, Figure 4d illustrates some natural frequency maps of individuals that did not match themselves.

## 4. Discussion

The present work aimed to adapt the methodology to map the brain’s natural frequencies (Capilla et al., 2022) from the group to the individual level. To improve the quality of the single-subject maps, we introduced two modifications to the original formulation of the algorithm. First, after performing k-means clustering on the entire group, a larger number of power spectra (∼1000) were assigned to each cluster centroid at the single-subject level. Second, we applied smoothing to the individual brain maps of natural frequencies. To assess the quality of the single-subject maps, we employed a brain fingerprinting approach. The underlying rationale was that, if individual brain patterns of natural frequencies are robust, they could be used to identify a target participant among others. Brain fingerprinting was conducted both within and between-sessions, achieving a high degree of accuracy in individual identification. Moreover, natural frequency fingerprints proved to be stable over time, even for MEG sessions conducted more than four years apart. Taken together, these results demonstrate that the brain’s natural frequencies can be reliably estimated at the single-subject level.

Our fingerprinting analysis showed an accuracy of nearly 93% for recognizing a target participant from two halves of the same recording. Accuracy dropped to 77% when the brain maps of natural frequencies came from two separate recordings. The lower accuracy values observed in the between-session group might be due to changes in brain state between the two sessions, although our results demonstrate that the drop in correct identifications is not systematically related with the passage of time. In addition, this could be explained by slight differences in co-registration between the first and second MEG sessions. Unlike previous MEG studies, we provided results at a voxel-by-voxel level. This substantially enhances the spatial resolution of the individual brain maps of natural frequencies and avoids the use of predefined anatomical parcellations, though at the expense of increased sensitivity to co-registration.

Overall, our identification accuracy results are in line with those found in related research. For example, in their seminal work, Finn et al. (2015) used individual functional connectivity profiles derived from resting fMRI data recorded on two consecutive days to identify participants, achieving a success rate of over 92%. MEG studies show more variable results. For example, Colenbier et al. (2023) attained modest identification accuracy of approximately 60% when attempting to identify individuals based on the spectral power of resting-state oscillatory activity, although identifiability raised up to 100% when participants were engaged in explicit tasks. Da Silva Castanheira et al. (2021) achieved success rates of around 95% using both spectral and functional connectivity features within the same MEG session. When comparing participants who underwent a second MEG session with a gap of up to 2.8 years, identification accuracy for functional connectomes decreased slightly to 89%, whereas spectral fingerprinting remained robust, with higher accuracy scores of nearly 98%.

Identifiability and differentiability are measures of the reliability in identifying a participant within the cohort and their individual saliency, respectively. Our identifiability results of about 17% and 32% for within- and between-session fingerprinting and differentiability of 2.03-3.26 are also comparable with findings from previous MEG studies (Colenbier et al., 2023; Da Silva Castanheira et al., 2021; Sareen et al., 2021). Importantly, we did not observe any significant correlation between identifiability and the number of days between MEG sessions, in agreement with Da Silva Castanheira et al. (2021). This implies that intrinsic oscillatory activity is an inherent and stable feature of brain function, unique to each individual. It would be interesting to explore the temporal boundaries of this stability in future research, as the longest interval between the two sessions in our data was 4.6 years.

Our study brings forward two important innovations in the field of MEG fingerprinting. First, we introduced a new feature of electrophysiological brain activity that can be used to recognize an individual among others, i.e., the natural frequency. Previous research has used spectral power or functional connectivity measures, based on either amplitude or phase (Colenbier et al., 2023; Da Silva Castanheira et al., 2021, 2024; Sareen et al., 2021). Here, we have demonstrated that the typical oscillatory frequency of each brain region is also a stable feature that can be employed to identify individuals. Importantly, the algorithm to obtain the brain’s natural frequencies provides precise values, making the use of frequency bands unnecessary (Capilla et al., 2022). Frequency band boundaries are usually established based on conventional criteria, rather than reflecting the brain’s natural oscillatory patterns, which might lead to misclassification of similar oscillations across species, life stages, or even different states within the same individual (Boersma et al., 2011; Buzsáki, 2006; Buzsáki et al., 2013). Additionally, the brain pattern of natural frequencies is a relatively simple index of brain function, since it involves only one value per voxel (1925 elements in this study, compared to the up to 30,000 used in the spectral fingerprinting of previous research) (e.g., Da Silva Castanheira et al., 2021). Although this is not an advantage in itself, it indicates that a simple metric can yield good identification results, which could be combined with others to further improve identification (e.g., natural frequency along with mean spectral power at the natural frequency).

The second innovation is the voxel-level resolution of our brain maps. Previous studies have often parcellated the brain into regions of interest to reduce the number of features used for fingerprinting (e.g., 68 regions in Da Silva Castanheira et al., 2021, or 148 regions in Sareen et al., 2021). However, parcellation is not trivial when applied to brain fingerprinting. Finn et al. (2015) showed that, in terms of identification rates, network definitions based on high-resolution parcellation (268 regions) outperformed those based on fewer regions, which may average out individual variability. Moreover, Capilla et al. (2022) found that the regional organization of intrinsic oscillatory brain activity does not completely overlap with typical anatomical parcellations, as some abrupt transitions in natural frequencies lack anatomical correspondence. Therefore, parcellation, if used, should rely on functional rather than anatomical criteria. Future research could develop a functional parcellation based on the brain’s intrinsic oscillatory activity, mirroring the parcellation of the brain derived from the resting-state fMRI functional connectome (Eickhoff et al., 2018; Thomas Yeo et al., 2011).

In conclusion, this study demonstrates that it is possible to obtain single-subject brain maps of natural frequencies that are robust enough to reliably identify a specific individual. The in-depth analysis of oscillatory frequencies adds a new layer to the more commonly studied power and phase, potentially offering valuable insights into the brain’s communication code. Furthermore, the ability to generate brain maps of natural frequencies at the individual level represents a significant advancement, as access to single-subject data is essential for conducting statistical analyses. This approach may also pave the way for detecting patterns of intrinsic oscillatory activity that deviate from normative patterns and could be indicative of particular pathologies, the so-called oscillopathies (Buzsáki et al., 2013).

## Author contributions

Lydia Arana: Conceptualization, Data curation, Formal analysis, Methodology, Software, Visualization, Writing – original draft preparation; Juan José Herrera-Morueco: Methodology, Writing – review & editing; Javier Santonja: Conceptualization, Resources, Writing – review & editing; Almudena Capilla: Conceptualization, Data curation, Funding acquisition, Methodology, Writing – original draft preparation.

## Declaration of competing interest

Lydia Arana is employed by CONECTIVA BRAIN HEALTH S.L. while pursuing a PhD at Universidad Autónoma de Madrid; however, this work is independent of CONECTIVA BRAIN HEALTH S.L, and no funding, data, or resources from the company were used in the study. The authors declare no competing interests related to this manuscript.

## Acknowledgements

We are thankful to the researchers and technicians involved in The Open MEG Archive (OMEGA) project for recording and making publicly available the database employed in the present study. The authors also thank Enrique Stern and Jorge San Segundo for valuable advice regarding the manuscript.

This work was supported by Ministerio de Ciencia e Innovación/Agencia Estatal de Investigación, Spain/FEDER/FSE+, UE (MCIN/AEI/ 10.13039/501100011033/FEDER/FSE+, UE; PID2021-125841NB-I00 to AC); and the Comunidad de Madrid, Spain (IND2022/SOC-23652 to LA).

